# The influence of phosphatidylserine localisation and lipid phase on membrane remodelling by the ESCRT-II/ESCRT-III complex

**DOI:** 10.1101/2020.04.21.053389

**Authors:** Andrew Booth, Christopher J. Marklew, Barbara Ciani, Paul A. Beales

## Abstract

The endosomal sorting complex required for transport (ESCRT) organises in supramolecular structures on the surface of lipid bilayers to drive membrane invagination and scission of intraluminal vesicles (ILVs), a process also controlled by membrane mechanics. However, ESCRT association with the membrane is also mediated by electrostatic interactions with anionic phospholipids. Phospholipid distribution within natural biomembranes is inhomogeneous due to, for example, the formation of lipid rafts and curvature-driven lipid sorting. Here, we have used phase-separated giant unilamellar vesicles (GUVs) to investigate the link between phosphatidylserine (PS)-rich lipid domains and ESCRT activity. We employ GUVs composed of phase separating lipid mixtures, where unsaturated DOPS and saturated DPPS lipids are incorporated individually or simultaneously to enhance PS localisation in liquid disordered (L_d_) and/or liquid ordered (L_o_) domains, respectively. PS partitioning between the coexisting phases is confirmed by a fluorescent Annexin V probe. Ultimately, we find that ILV generation promoted by ESCRTs is significantly enhanced when PS lipids localise within L_d_ domains. However, the ILVs that form are rich in L_o_ lipids. We interpret this surprising observation as preferential recruitment of the L_o_ phase beneath the ESCRT complex due to its increased rigidity, where the L_d_ phase is favoured in the neck of the resultant buds to facilitate the high membrane curvature in these regions of the membrane during the ILV formation process. L_d_ domains offer lower resistance to membrane bending, demonstrating a mechanism by which the composition and mechanics of membranes can be coupled to regulate the location and efficiency of ESCRT activity.

## Introduction

The precise molecular composition of lipid membranes determines the physicochemical properties that regulate biological cell homeostasis^1^, such as membrane fluidity and phase behaviour. Lipid membrane fluidity and phase behaviour influences the mechanical properties of membranes (e.g., bending modulus^2^, spontaneous curvature^3^) and phase separation imparts spatial ordering of lipids, which can play a role in biological signalling (in so-called ‘lipid rafts’).^4,5^ This spatial ordering of lipids modulates the local mechanical properties of membranes^2,6^ which plays a key role in signalling mechanisms comprising of membrane fusion and fission events.^4,5,7^

Membrane tension regulates the activity of membrane-remodelling protein complexes such as dynamin^8^ and the Endosomal Sorting Complex Required for Transport (ESCRT), which drives membrane budding and scission in multivesicular body biogenesis, viral budding, repair of plasma, nuclear and organelle membranes.^9–11^ The core scission machinery of ESCRT is comprised of the ESCRT-III complex and the adaptor complex ESCRT-II, which binds specifically to phosphatidylinositoside phosphates (PIPs) and ESCRT-III initiates membrane invagination.^12^ In budding yeast, ESCRT-III assembles from four key subunits (Vps20, Snf7, Vps24 and Vps2) and the AAA+ ATPase Vps4 that regulates the formation of ESCRT-III three-dimensional spiral structures.^13^ These structures are capable of stabilising membrane buds and constricting membrane necks in an ATP-dependent manner.^13^

The budding and scission activity of the ESCRT-II and ESCRT-III complexes can be reconstituted *in vitro* by following the generation of intralumenal vesicles (ILVs) within giant unilamellar vesicles (GUVs) after the addition of purified ESCRT components.^12,14^ In this bulk-phase encapsulation assay, only ESCRT-generated ILV compartments contain fluorescent extravesicular medium in their lumens allowing the quantification of ESCRT-mediated membrane-remodelling upon a change in experimental conditions. *In vitro* systems to study how bulk phase is encapsulated into membrane compartment by protein complexes such as ESCRT also represent a useful biomimetic tool to create multi-compartment architectures incorporating spatially segregated, chemically distinct environments and possess significant promise in the construction of complex artificial cells.^15,16^

It is well established that specific lipid headgroups play key roles in signalling and protein recruitment processes.^17^ Anionic PIPs and phosphatidylserine (PS) lipids are necessary to initiate ESCRT activity at the membrane.^18^ Specifically, PS lipids are necessary for the binding of ESCRT-III subunits prior to complex assembly and membrane remodelling^9,19–21^ whereas ESCRT-II subunits specifically binds PI(3)P^22^ though phosphoinositides are not strictly required for scission activity.^14^

The affinity of ESCRTs for specific lipids therefore provides an opportunity to employ phase separated model membranes to investigate how spatially distinct lipid distribution between membrane phases of differing fluidity and mechanics influences their function and activity *in vitro*. Simple mixtures of saturated lipids, unsaturated lipids and cholesterol can be designed to phase separate into coexisting fluid membrane phases that are thought to mimic the structural properties of native lipid rafts.^23–26^ The liquid disordered (L_d_) phase is enriched in unsaturated lipids and is characterised by higher fluidity and lower bending rigidity, while cholesterol and saturated lipids concentrate in more viscous and rigid liquid ordered (L_o_) domains. Equilibration of phase separating GUV membranes results in membrane textures that coarsen into large microscale domains that minimise the interfacial tension between coexisting phases. These domains can easily be visualised by fluorescence microscopy techniques due to preferential partitioning of lipophilic trace fluorophores between distinct membrane phases.^27,28^

Here we show that ESCRT-II/ESCRT-III activity is increased in PS-rich L_d_ microdomains with the newly formed ILVs containing predominantly L_o_ phase membranes. Given the relationship between mechanical properties and membrane phases, our data suggests further insight into the mechanism of mechanical deformation of membranes by the ESCRT complex. Furthermore, if specific lipids can indeed be used to ‘target’ ESCRT-driven ILV formation to membrane microdomains then we can envisage strategies for generating ILVs with physicochemical properties selectively distinct from their parent membrane.

Our data provides further insight into the *in vitro* mechanism of mechanical deformation of membranes by the ESCRT complex and might aid the understanding of ESCRT-related events at biological membranes.

## Results and discussion

### Strategy

We and others^14,15,29^ have successfully used GUVs composed of homogeneous PC:Cholesterol:PS mixtures to reconstitute the membrane remodelling activity of the ESCRT-II/ESCRT-III complex. Based on phase diagrams available in the literature for DOPC:DPPC:chol mixtures,^30^ we selected a baseline membrane composition of 35:35:30 as it is fairly central within the liquid ordered (Lo) - liquid disordered (Ld) coexistence region of the phase diagram. 13 mol% of PC lipids were exchanged for the corresponding PS lipid with the same acyl tails (e.g.: DOPC for DOPS) in each formulation, with either 13 mol% DOPS, 13 mol% DPPS or 6.5 mol% DOPS + 6.5 mol% DPPS. **Figure 1** depicts the expected distribution of PS lipids in our formulations due to preferential partitioning of these lipids between the coexisting phases and the hypothesised resulting ILV membrane compositions.

**Figure 1:**
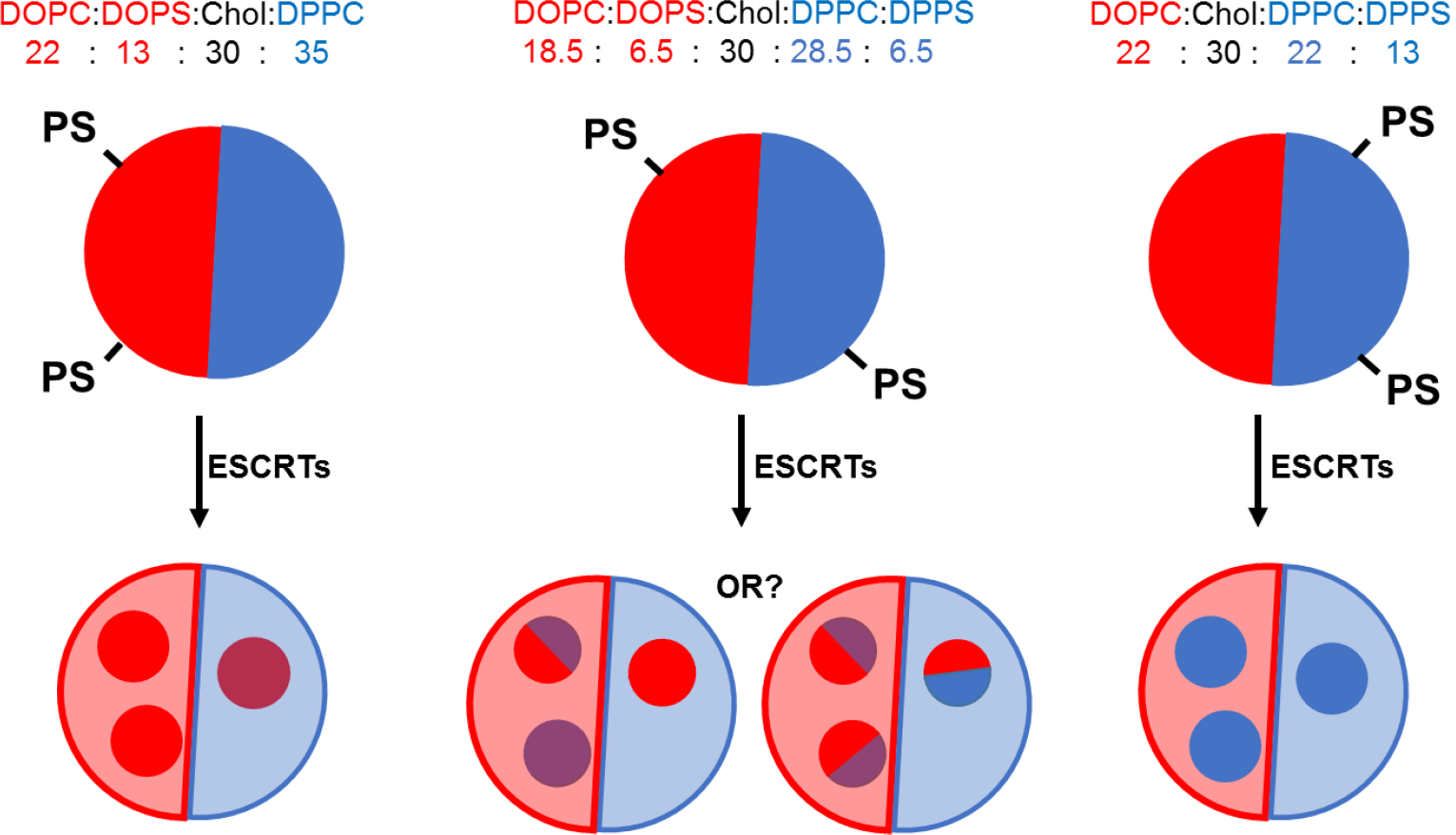
Schematic of hypothesised outcomes from ESCRT-driven ILV formation activity against GUVs with differently distributed PS-headgroup lipids. ILV membrane composition may reflect the relative distribution of the phase(s) with increased PS concentration. Red: L_d_ membrane, Blue: L_o_ membrane.

### Controlling the phase distribution of PS-headgroup lipids

Confocal microscopy of GUVs treated with the PS-binding protein Annexin V, fluorescently labelled with AlexaFluor488 (AF-488-AxV), was used to assess PS localisation in each formulation with respect to membrane phase domain (**Figure 2**). Phase character was indicated by the partitioning of Naphtho[2,3-a]pyrene (Np) into L_o_ domains and of 1,2-dioleoyl-sn-glycero-3-phosphoethanolamine-N-(Lissamine rhodamine B sulfonyl) ammonium salt, (‘Rh-PE’) into L_d_ domains.^27^ The great majority of GUVs were observed to have roughly hemispherical L_o_/L_d_ distributions, although some striped and spotted domains were also sporadically observed The expected phase-enriched distributions of PS lipids were obtained reliably, (L_o_ PS enrichment with DPPS, L_d_ enrichment with DOPS, and comparable distributions in the 6.5 mol% DPPS, 6.5 mol% DOPS ‘1:1’ mixture. The distribution of AF-488-AxV within enriched phases was highly uniform other than some isolated cases where enhanced AF-488-AxV fluorescence was observed along phase boundaries in the ‘1:1’ PS composition, but this was rare.

**Figure 2:**
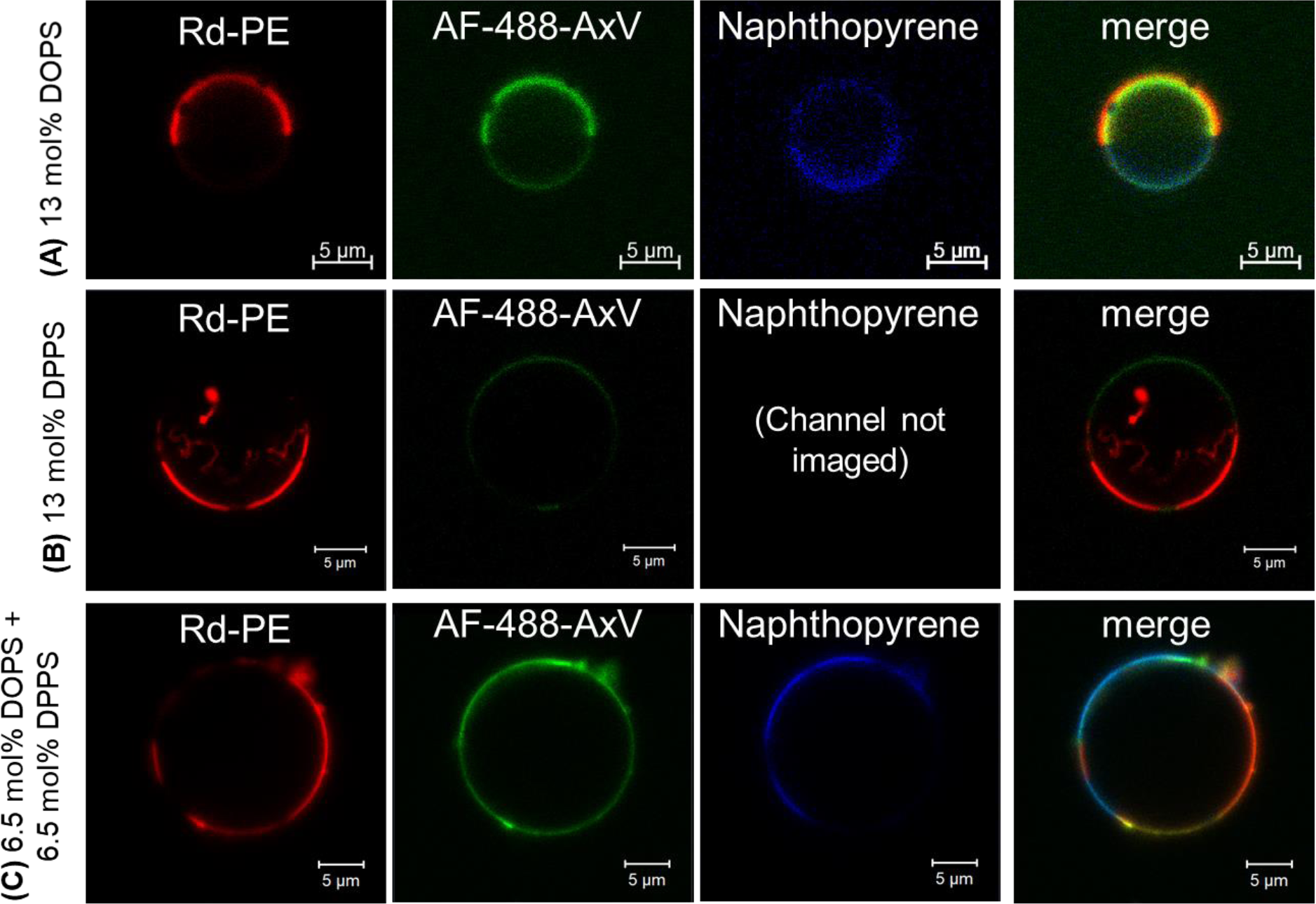
Confocal microscopy of phase separated GUVs. **(A)** 28.5 : 28.5 : 13 : 30 DPPC : DOPC :DOPS : Chol, **(B)** 28.5 : 28.5 : 13 : 30 DPPC : DOPC :DPPS : Chol, **(C)** 28.5 : 28.5 : 6.5 : 6.5 : 30 DPPC : DOPC :DOPS : DPPS : Chol. Rh-PE preferentially partitions into L_d_ domains, Napthopyrene (Np) preferentially partitions into L_o_ domains, AF488-AxV selectively binds to PS lipids to indicate distribution with respect to the domains of different membrane phases.

### PS in the L_d_ phase enhances efficiency of ILV formation

ESCRT activity was assessed by ILV counting. Membrane-impermeable fluorescent dyes were added to GUV suspensions, followed by ESCRT proteins (50 nM Snf7, 10 nM each of ESCRT-II, Vps20, Vps24, Vps2 and Vps4, and 10 μM ATP.MgCl_2_), all added simultaneously. The presence of fluorescent dye in the lumens of free-floating ILVs, after incubation with ESCRT proteins, being a strong indication that those ILVs were formed as a result of ESCRT activity.^15^ The number of dye-containing ILVs was counted and then expressed as a number of ILVs per GUV volume equivalent, by taking the luminal volume observed during large area tile-scan imaging of GUV samples (cross sectional area x section thickness), divided by the volume of a typical 20 μm diameter GUV **Figure 3**. Automation of ILV counting was made possible by using the calcein-Co^2+^ fluorescent dye-quencher pair in this assay, as fluorescence in the bulk medium could be quenched after the ESCRT incubation period by addition of the membrane-impermeable quencher (CoCl_2_). This means that only ILVs containing calcein fluorescence had formed during the period of incubation with ESCRTs.

**Figure 3:**
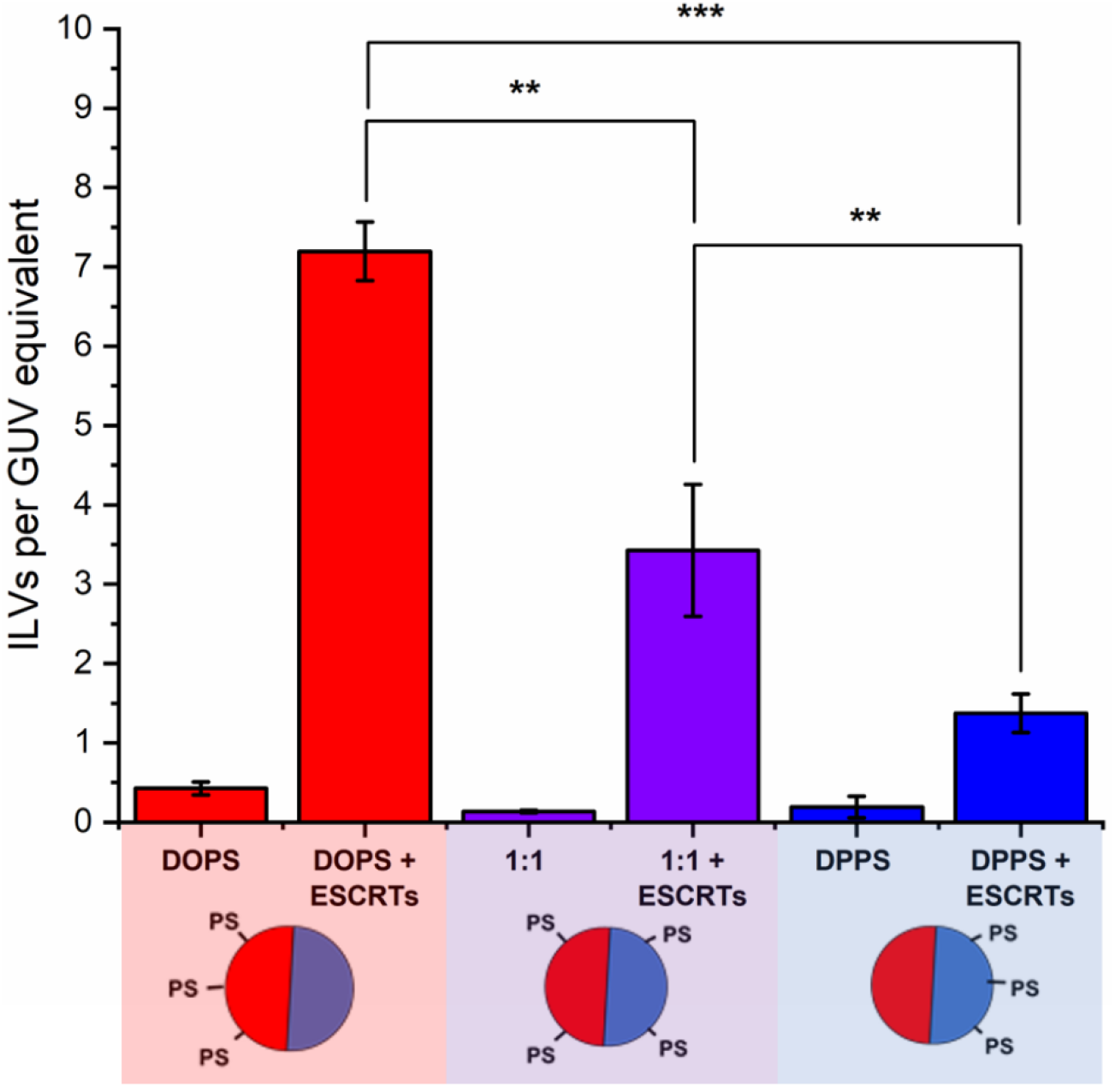
The impact of PS phase localisation on the ILV-forming activity of ESCRTs. The graph shows the number of ILVs per GUV volume equivalent for GUVs incubated with the ESCRT-II/III complex (Snf7 50 nM, ESCRT-II, Vps20, Vps24, Vps2, Vps4 (10 nM) and ATP.MgCl_2_ 1 μM. ILVs are determined to be formed post-ESCRT addition by the presence of calcein fluorescence. GUVs either contained DOPS localised to the L_d_ domains, DPPS localised to the L_o_ domains, or a 1:1 mix of DOPS and DPPS that fairly evenly distributed between the coexisting phases. **Diagram**: Red: L_d_ membrane, Blue: L_o_ membrane.

The results, shown in **Figure 3**, indicate that the relative localisation of PS-headgroup lipids in phase-separated GUVs has a significant influence on the extent of ESCRT-induced ILV formation. The highest ILV formation activity was seen within GUVs containing 13 mol% DOPS, where PS lipids are predominantly located in the L_d_ phase domains of the GUVs. In contrast, the lowest ILV formation activity was seen in GUVs where PS is preferentially located in the L_o_-phase (13 mol% DPPS GUVs). Finally, where PS is found in both phases (1:1) an intermediate degree of ILV formation was observed.

This suggests that the concentration of PS lipids in Ld domains also regulates the degree of ESCRT activity. We can rationalise these findings by taking into consideration the lower bending rigidity of each phase; the lower bending rigidity of an Ld phase, offers lower mechanical resistance to ESCRT-induced membrane deformation than a Lo domain.^2,31^ ESCRT activity is negatively regulated by membrane tension, in a manner that is likely analogous to suppression of activity by higher bending rigidity.^15^ We conclude that ESCRT activity must be higher at PS-containing L_d_ domains where the membrane is more susceptible to deformation.

### ESCRT-induced ILVs consist of L_o_ phase membranes

Surprisingly, the ILVs that form mostly consist of L_o_ membranes, despite localisation of PS lipids in L_d_ domains (**Figure 4**, blue bars). This is determined by the observed depletion of the L_d_-partitioning Rh-PE probe from the membranes of ILVs, indicating a lack of liquid disordered phase membrane. In contrast with the few ILVs present in the GUV lumen prior to addition of proteins, lacking encapsulated calcein, and formed as a random by-product of the electroformation process, (**Figure 4**, red bars), showed greater L_d_ compared to those formed by ESCRT activity.

**Figure 4:**
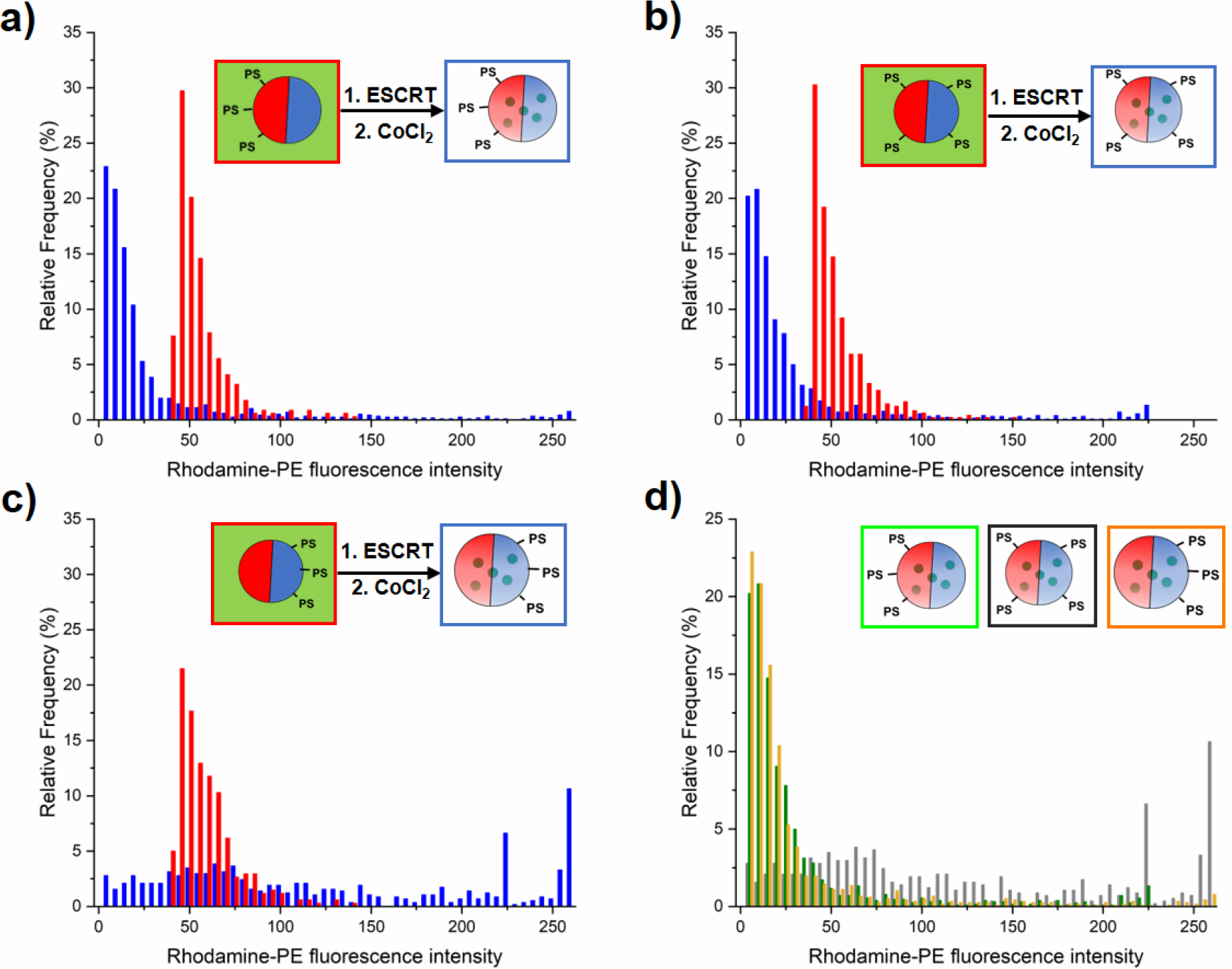
ILV membranes are predominantly Lo phase. Histograms of mean Rhodamine-PE fluorescence intensity in ILVs, a measure of the phase character of the membranes. **Red plots**: Intrinsic ILVs (ESCRT-independent). **Blue plots**: ESCRT-induced ILVs. **a)** 13 mol% DOPS GUVs, **b)** 6.5 mol% DOPS, 6.5 mol% DPPS GUVs, **c)** 13 mol% DPPS GUVs, **d)** overlaid post-ESCRT ILV data for each composition (green = DOPS, grey = 1:1 mixture, yellow = DPPS). **Diagrams**: Red: L_d_ membrane, Blue: L_o_ membrane.

We therefore quantify the relative phase character of ILVs formed by ESCRTs and those that occur intrinsically to ascertain the significance of this observation. We quantified the area mean fluorescence intensity of the Rh-PE for each individual ILV present before and after the addition of ESCRT proteins, taking this as a measure of the proportion of the L_d_ phase in the volume of the ILV (**Figure 4**). Analysis of Rh-PE fluorescence is more robust than analysis of the Np fluorescence, due to the weaker overall fluorescence and its minor overlap with the fluorescence emission of encapsulated calcein.

The 13 % DOPS (**Figure 4.a**) and 1:1 DOPS:DPPS (**Figure 4.b**) GUV samples, show a distinctly lower mean Rh-PE fluorescence than intrinsic ILVs, We interpret this as a decrease in L_d_ character of the membrane, or alternatively, a significant enrichment of L_o_ phase in ESCRT-induced ILVs.

This observation was surprising since it would be reasonable to anticipate that the ILVs form from the PS-enriched phase of the parent GUV membrane and hence have a similar composition and phase character. However, this result is consistent with previous suggestions of localisation of ESCRT complexes at phase boundaries, where local membrane curvature may help to nucleate the assembly of ESCRT-III.^18^

The GUVs containing 13 mol% DPPS exhibit ILVs with a very broad range of mean fluorescence intensities with no obvious compositional preference (**Figure 4. C**). This lack of selectivity for L_o_ or L_d_ domains and significantly reduced ILV formation activity (**Figure 3**) suggests ESCRT activity is dysregulated when PS is enriched in the L_o_ domains, the tighter packing of lipids in this phase may inhibit membrane insertion of the amphipathic helix of Snf7, thereby restricting its normal assembly and function.^32,33^

A representative image of DOPS-containing GUVs after incubation with ESCRTs is shown in **Figure 5 A**. Here, the significant reduction in Rh-PE fluorescence in ESCRT-induced (those containing green calcein fluorescence) relative to intrinsic ILVs is strikingly illustrated. However, **Figure 5 A** also illustrates a common observation in GUV samples with compositions containing DOPS (13 mol% DOPS, 1:1 DOPS:DPPS), that GUVs with a high abundance of ESCRT-induced ILVs tend to have a predominantly L_o_ parent membrane. This is a highly counterintuitive finding as the expectation would be that any mechanism that favours the formation of L_o_ ILVs necessarily leads to the removal of L_o_ lipids from the parent GUV membrane. Having already shown that PS-localisation in the L_o_ phase inhibits ESCRT activity, it is unlikely that ESCRTs have preference for L_o_-only vesicles in the DOPS GUV samples. Indeed L_o_-only GUVs were not observed prior to ESCRT addition, suggesting that such vesicles are ESCRT-induced and high ILV formation activity in a single vesicle likely results in shedding of L_d_ phase from the parent membrane. We will show evidence for this in rare observations of stalled ESCRT-induced membrane buds.

**Figure 5:**
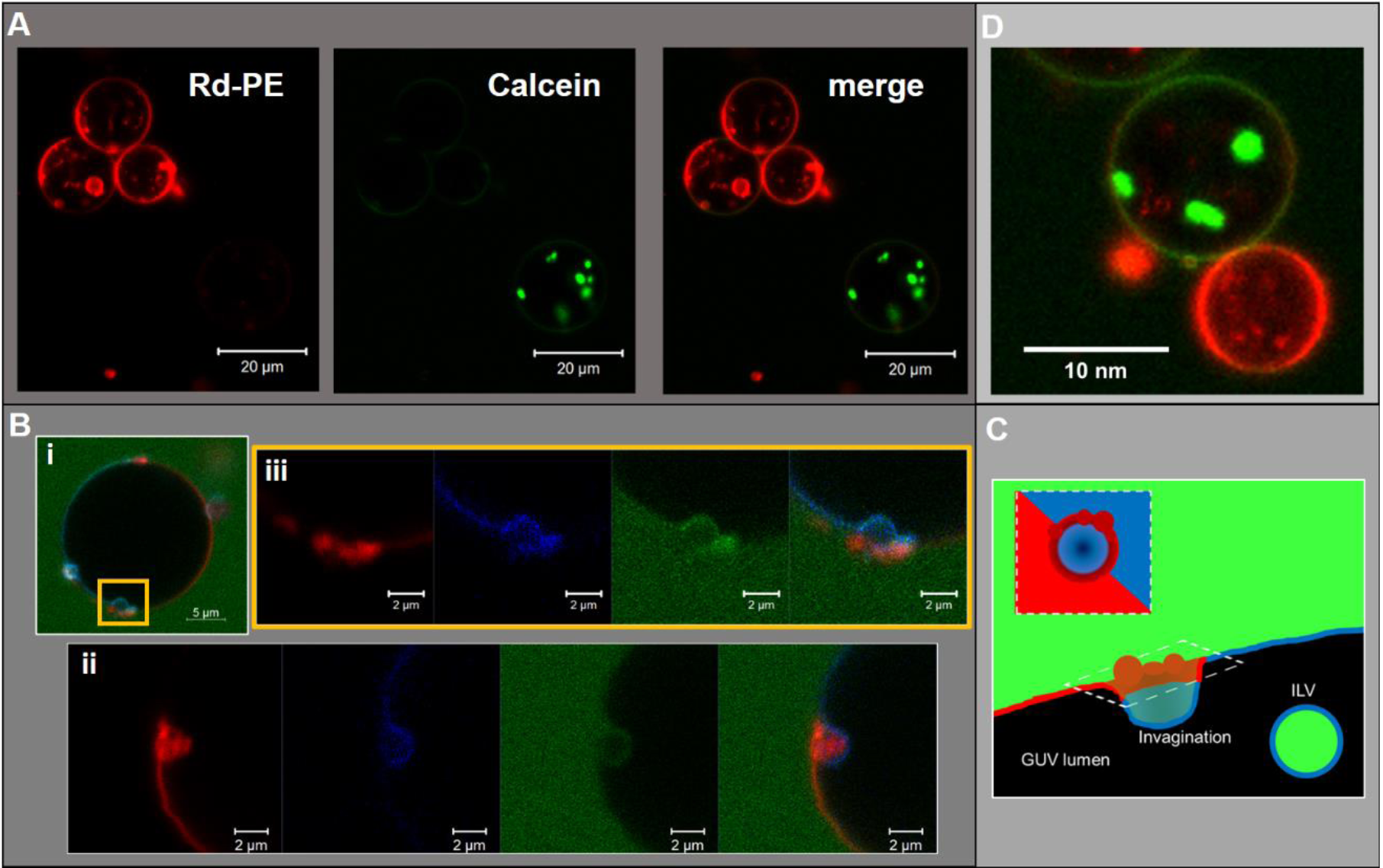
Representative images of observed phase characteristics in GUVs incubated with ESCRTs. **A**: High yields of ESCRT-induced ILVs in an individual GUV causes depletion of L_d_ phase from the parent membrane. Confocal microscopy of GUVs (13 mol% DOPS) after incubation with ESCRT proteins and quenching of background calcein fluorescence with CoCl_2_. Left: Rh-PE, Centre: calcein, Right: merge. **B**: Confocal microscopy images of GUVs with apparent stalled ILV buds. **i** and **ii** are distinct GUVs, **iii** is an enlargement of the yellow box in I, left to right; Rh-PE, Np, fluorescein labelled dextran, 70 kDa, both GUVs are ‘1:1’ composition (6.5 mol% each of DPPS and DOPS. **A(i)** is an enlargement of the area designated by the yellow square on **A Green**: fluorescein labelled dextran (70 kDa), **Red**: Rhodamine-PE, **Blue**: Np. **C:** Schematic interpretation of ILV bud phase character distribution based on images **5.B (i)** and **(ii). D:** L_o_ GUV with calcein containing ILVs with adherent L_d_ GUV, possibly a result of ESCRT-induced abscission.

### Stalled ILV ‘buds’ reveal insights into the mechanism of L_o_-enrichment in ILVs

We observed that a small number of GUVs with a 1:1 DOPS:DPPS composition contained stalled ILV buds (**Figure 5 B**). One mechanism by which these stalled ILV buds may appear is when the membrane tension of a particular GUV is sufficient to allow initial invagination of the ILV bud, but mechanical resistance stalls ILV formation prior to scission of the bud neck. We have previously observed a number of these stalled buds in homogeneous GUV membranes interacting with ESCRT proteins.^15^ Alternatively, rare assembly errors in the formation of these complexes may have resulted in incompetent ESCRT machinery that is unable to complete the final neck closure and scission steps.

These nascent buds are yet to be closed off from the extravesicular medium given that we observe a continuity of fluorescein-labelled dextran ([70 kDa]) from the bulk phase into the bud volume. These buds have a neck of approximately 2 μm diameter, which correlates with the typical size range of ILVs observed in these experiments (∼1-2 μm diameter). The membrane phases show an unusual distribution in these bud structures where buds occur at or near to a phase boundary, with the portion of the membrane that buds into the parent GUV lumen being predominantly L_o_ (**Figure 5 B i)**. However, there is an accumulation of L_d_ membrane on the exterior of the parent membrane adjacent to the bud neck, which could be interpreted as a ‘collar’ of L_d_ membrane (see **Figure 5 C**). This contrasts somewhat with the structure observed in **Figure 5 B ii**, where again, a L_o_ bud is observed at a phase boundary, with L_d_ accumulation around the bud neck, but there is also a protrusion of L_d_ material into the bud lumen. Despite the infrequent observation of this phenomenon, there is a clear correlation with the prevalence of L_o_ phase character in ESCRT-induced ILVs as shown in **Figure 4**.

### Proposed mechanism of L_o_-enriched ILV formation

ILV formation is most efficient when PS lipids are localised within L_d_ domains of phase separated GUVs. Electrostatic association of the ESCRT-III proteins with PS lipids would be expected to concentrate initial binding onto L_d_ domains. Curvature generated at phase boundaries^34^ may also preferentially nucleate ESCRT-III assembly at these sites.^18^ Assembly of ESCRT complexes on the membrane likely increases the local membrane rigidity, suppressing thermal undulations. Suppression of membrane undulations in adhesion plaques has previously been shown to promote formation and localisation of more rigid membrane phases,^35^ and similar observations have recently been reported *in vivo*.^36^ Moreover, induced formation of L_o_ membrane beneath ESCRT-II assemblies has previously been reported on solid-supported lipid bilayers.^37^

Induced recruitment of L_o_ membrane beneath an ESCRT-II complex will create a line tension with the L_d_ phase at the domain boundaries. The energy cost of this phase boundary can be minimised by bending the membrane such that the circumference of the phase boundary decreases, promoting budding of the L_o_ domain.^38^ The L_d_ phase at the L_o_ domain boundaries is likely required to facilitate the high curvature in the bud neck, while the line tension within the L_d_-L_o_ phase boundary within the bud neck could enhance the efficiency of scission of the bud from the parent membrane, forming the ILV.^39–41^ It should be noted that the curvature of the L_o_ domain bud (radius of curvature ∼ 1 μm) is much lower than in the bud neck (radius of curvature ∼ 10s nm). These ESCRT-induced L_o_ buds may preferentially migrate to the macroscopic phase boundaries within the GUV, or preferentially nucleate in these regions of the vesicle, as observed in **Figure 5 B i, ii. Figure 5B** also shows strong evidence for involvement of L_d_ phase lipids in the neck of L_o_-rich membrane buds.

Notably, there is evidence for lipid sorting during vesicle budding processes *in vivo*, in the generation of endosomes, multivesicular bodies and exosomes.^42^ Interestingly, many lipidomics studies have reported enrichment of sphingomelin and cholesterol in endosomes and exosomes relative to their parent membrane (the plasma membrane), suggestive of a greater L_o_ phase character.^43–46^ The parallels with the L_o_-enrichment in phase separated GUV model membranes reported here are striking.

Importantly, ILV formation is strongly suppressed when PS lipids are enriched in L_o_ domains (**Figure 3**). Therefore, we propose that ESCRT-II/ESCRT-III, preferentially recruited to these L_o_ domains through electrostatic association with DPPS, have suppressed functional activity at these membrane domains. This conclusion is supported by the following considerations. In the *in vivo* system, the anchoring of ESCRT-III via membrane-inserting motifs (myristoyl moiety in Vps20^47^ or the N-terminal amphipathic helix in Snf7^33^) may be inhibited in the L_o_ phase, due to more ordered lipid packing and lower incidence of packing defects, resulting in lower surface concentration of bound protein.^48^ Furthermore, the high membrane curvature required in the neck of nascent buds would have a much higher energy cost for L_o_ phase membrane than L_d_ phase membrane, where the energy required to bend the membrane is a factor of ∼5-6 greater.^49^ ESCRT assembly on the higher rigidity L_o_ membrane would also not favour recruitment of a different membrane phase beneath the complex, where line tension would no longer be generated to favour budding and scission as described above. These factors likely combine to strongly suppress ILV formation generated from ESCRT binding to L_o_ domains, where the few ILVs that do form are highly dysregulated in membrane composition as observed in **Figure 4 C,D**.

The observed depletion of L_d_ lipids from GUV membranes induced by ESCRT-driven ILV formation is harder to conclusively explain due to limited experimental evidence. We present hypotheses in **Figure 6 B, C**. In **Figure 6 B** some of the dense L_d_ phase membrane accumulation around ILV bud necks (as seen in **Figure 5B**) is detached from the parent GUV membrane during the bud scission process. Following numerous ILV formation events, the L_d_ phase may be progressively depleted. Another source for L_o_-rich GUVs may be ESCRT-driven abscission of entire L_d_ domains from GUVs, generating separate L_d_-rich vesicles next to the L_o_-rich vesicle as depicted in **Figure 6C**. This is analogous to the role of ESCRT in cytokinetic abscission.^50^ A small number of GUVs were observed in this configuration in samples incubated with ESCRTs, a representative example is shown in **Figure 5 D**, but we currently have insufficient data to make any strong claims on the prevalence of this speculative mechanism. However, shedding of some membrane from GUVs caused by ESCRT activity should not be seen as too surprising since excess membrane shedding is required for the fission mechanisms at the basis of ESCRT activity *in vivo*.^51,52^

**Figure 6:**
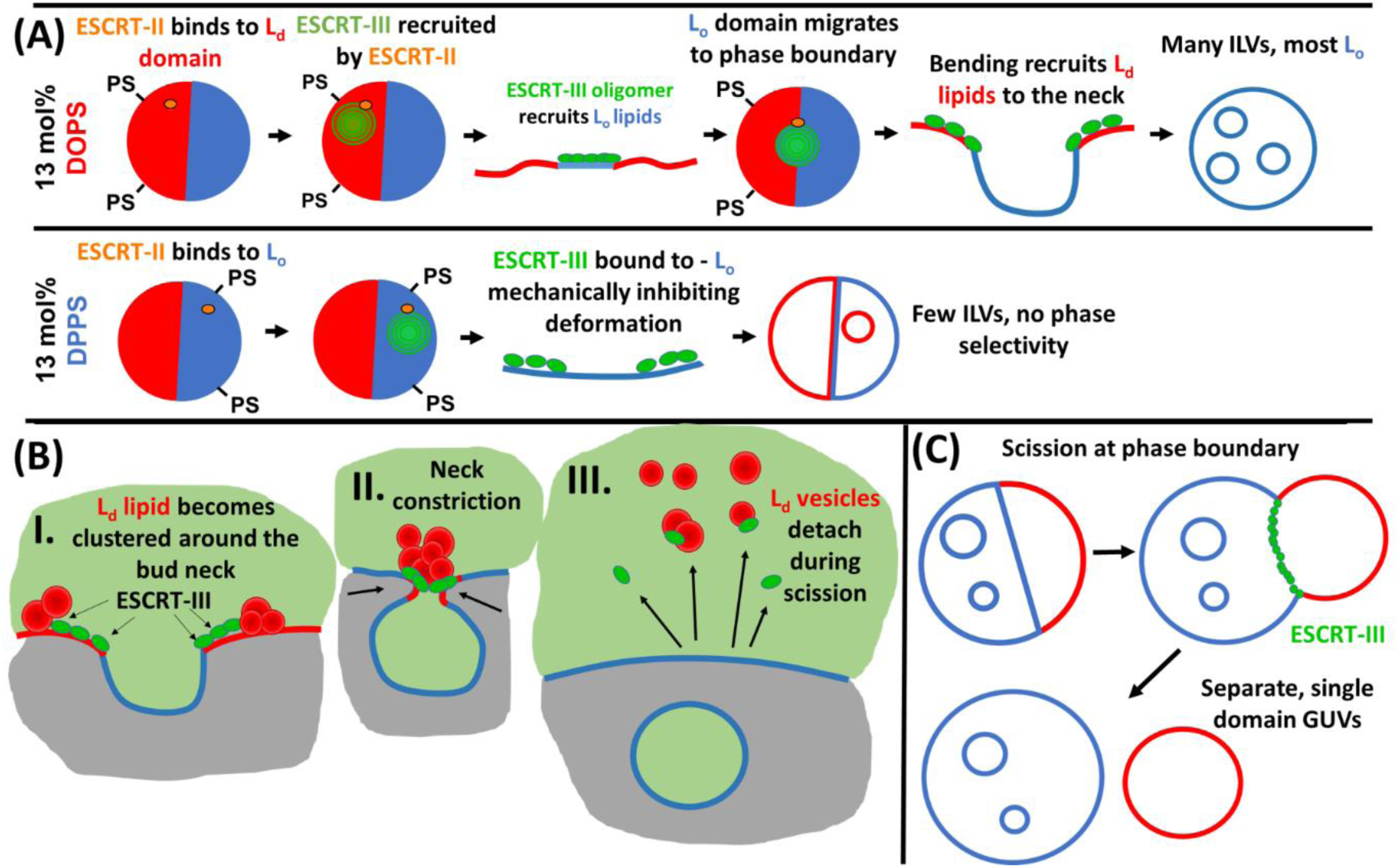
Proposed mechanistic models of the phase behaviour observed in ESCRT-induced ILV formation in phase separated GUVs. **A**: Model of ESCRT assembly and membrane remodelling activity in GUVs with PS lipids concentrated in L_d_ (13 mol% DOPS) or L_o_ (13 mol% DPPS) domains. **B**: Depletion of GUV membrane L_d_ domain by ejection of clustered L_d_ vesicles from the ILV bud neck during scission. **C**: Possible ESCRT-driven scission at GUV phase boundaries leading to distinct enriched or single domain GUVs as observed.

## Conclusions

We have shown that ESCRT-driven membrane remodelling in phase separated model membranes is most efficient when PS lipids are preferentially located in the less rigid L_d_ domains. At first sight this result is intuitive since these domains offer lower mechanically resistance to the budding deformations required for ILV formation. However, surprisingly the ILVs that form are strongly enriched in L_o_ membrane. Combining rare observations of stalled ILV formation events and membrane biophysical concepts, we propose a mechanism by which this L_o_ phase enrichment occurs. Generation of ILV proto-organelles with a different membrane composition to the GUV parent membrane may have a useful role in generating artificial cells that mimic the variability of lipid composition between organelles and the plasma membrane of natural cells. Furthermore, the comparison of our observations with the enrichment of L_o_-preferring lipids in endosomes and exosomes is compelling, suggestive of a similar phase-enrichment potentially occurring for ESCRT-generated ILVs *in vivo*.

## Materials and methods

### Materials

All lipids; DOPC (1,2-dioleoyl-sn-glycero-3-phosphocholine), DOPS (1,2-dioleoyl-sn-glycero-3-phospho-L-serine (sodium salt)), DPPC (1,2-dipalmitoyl-sn-glycero-3-phosphocholine), DPPS (1,2-dipalmitoyl-sn-glycero-3-phospho-L-serine (sodium salt)), Cholesterol (ovine) and Rh-PE (1,2-dioleoyl-sn-glycero-3-phosphoethanolamine-N-(lissamine rhodamine B sulfonyl) (ammonium salt)) were obtained from Avanti Polar Lipids Inc. (Alabaster, AL, USA). Np (Naphtho[2,3-a]pyrene) was obtained from Tokyo Chemical Industries UK Ltd. Calcein, NaCl, Tris buffer, CoCl_2_ (hexahydrate), MgCl_2_, sucrose, ATP (Adenosine 5′-triphosphate disodium salt hydrate) were all obtained from Sigma-Aldrich Company Ltd. (UK). 8-chamber glass-bottomed microscope slides (Cat. No. 80827) were used for all confocal microscopy and were obtained from ibidi GmbH. Annexin V AF488 conjugate (Cat. No. A13201) was obtained from Thermo Fisher Scientific Inc. All proteins were expressed and purified in-house.

### Protein expression

The plasmids for overexpression of *Saccharomyces cerevisiae* ESCRT-II (Vps22, Vps25, Vps36) and ESCRT-III subunits Vps20, Snf7, Vps2 and Vps4 are a gift from James Hurley (addgene plasmids #17633, #21490, #21492, #21494, #21495).^14,53^ The *Saccharomyces cerevisiae* Vps24 gene was subcloned into a pRSET(A) expression vector, containing a His6-tag and Enterokinase protease cleavage site. We have previously reported the detailed expression and purification methods for the ESCRT-II and ESCRT-III proteins used in this the study, including characterisation and quality control data in reference.^15^

### Electroformation of GUVs

Giant unilamellar vesicles (GUVs) were prepared using the electroformation method. Briefly, the desired lipid mixture is applied to indium tin-oxide coated glass slides (8-12 Ω/sq, Sigma-Aldrich), as a chloroform solution (15 μL, 0.7 mM) and dried under nitrogen to form a thin film. Lipid compositions were either: **‘13 mol% DOPS’**: 35 : 22 : 13 : 30 DPPC : DOPC :DOPS : Chol, **‘13 mol% DPPS’**: 22: 35 : 13 : 30 DPPC : DOPC :DPPS : Chol, or **‘1:1’** 28.5 : 28.5 : 6.5 : 6.5 : 30 DPPC : DOPC :DOPS : DPPS : Chol. An additional 0.5 mol% of Rh-PE and 1 mol% of Np were added to each stock solution.

Two slides, coated with the desired lipid mixture, were assembled into an electroformation chamber by separation of their conductive, lipid-coated surfaces by a silicone rubber gasket (∼2 mm thickness). Electrical contacts were made to each slide using copper tape and these were connected to a function generator. The chamber was then filled with sucrose solution (300 mM) and sealed with a silicone rubber plug. The chambers were then placed into an oven at 60 °C. An AC field was applied to the chambers; 0.1 V_peak-to-peak_, 10 Hz, sinusoidal waveform, ascending to 3 V_pp_ over 10 min, and maintained at 3 V for 2 h, after which the frequency of the oscillation was reduced from 10 to 0.1 Hz over 10 min. At this point the GUV-containing sucrose solution was harvested using a syringe.

Osmotic relaxation of membrane tension was then performed. GUVs were diluted 1:4, vol:vol, GUV suspension:hypertonic buffer, with 10 mM Tris buffered saline (∼150 mM NaCl; pH 7.4), adjusted to 10 mOsm kg^-1^ higher osmolarity than the sucrose solution and stored overnight at 4 °C prior to use. Solution osmolarity was measured using an Advanced Instruments 3320 osmometer.

### Confocal microscopy

All experiments were performed on Zeiss LSM 700 or 880 systems, imaging with a Plan-Apochromat 40x/1.4 Oil DIC M27 objective lens, NA = 1.4, 405 nm, 488 nm, and 561 nm lasers. GUV suspensions were placed in chamber slides (µslide, 8 well, glass bottomed, ibidi GmbH). Glass surfaces in the chamber slides were passivated prior to use by addition of 5 wt% bovine serum albumin solution in deionised water, incubation for 5 min, followed by copious rinsing with milliQ water and finally blown dry using a stream of nitrogen gas. 200 μL aliquots of GUV suspension were added to each well.

### Annexin V PS localisation assay

Prior to ESCRT assays, separate aliquots of GUVs from the same samples were treated with Annexin V, to confirm PS distribution was as expected. AlexaFluor488-labelled Annexin V (Thermo Fisher cat. No.: A13201) (2 μL per 200 μL GUV suspension) was added to desired wells and incubated for ∼5 min before imaging.

### ESCRT-induced ILV counting assay

Calcein (2 μL of a 4 mM stock solution in osmotically balanced saline buffer [10 mM Tris, pH 7.4]) was added to each well (200 μL GUV suspension), immediately followed by ESCRT proteins; ESCRT-II (Vps22, Vps25, Vps36), Vps20, Snf7, Vps24, Vps2 and Vps4 and ATP.MgCl_2_ were added to each desired GUV suspension (200 μL) to give a final concentration of 10 nM for ESCRT-II, Vps20, Vps24, Vps2, Vps4, 50 nM for Snf7, and 10 μM ATP.MgCl_2_. After a 20 min incubation period, CoCl_2_ (2 μL of a 4 mM stock solution in osmotically balanced saline buffer [10 mM Tris, pH 7.4]), was added to quench calcein fluorescence in the extravesicular solution. Alternatively, fluorescently labelled dextrans of varying molecular weights were added to assess uptake of larger species into ILV lumens. All imaging was performed with a pinhole setting that corresponded to a 3.1 μm section thickness, allowing for a greater volume of a GUV’s lumen to be captured and to increase the likelihood of ILVs being entirely captured within the visible z-dimension, to improve the accuracy of membrane phase characterisation of both ILVs and GUVs. Large area tile-scanning was performed in order to capture sections through as high a number of GUVs as possible, to allow statistical analysis across the GUV population. Images were analysed using the Zeiss Zen software and Fiji. To improve comparability between results, ILV counts were expressed as ‘ILVs per GUV volume equivalent’, being the total GUV lumen observed (cross-sectional area x section thickness), divided into units of typical 20 μm diameter spherical GUV volumes.

### ILV phase character analysis

The phase character of ESCRT-induced ILVs was assessed by calculating the mean fluorescence intensity of Rh-PE, over the area of the ILV-cross section, as a measure of the amount of L_d_ membrane present. ESCRT-induced ILVs were identified as those which had calcein fluorescence in their lumen. This was compared to the same analysis of intrinsic ILVs occurring in GUV samples prior to the addition of ESCRT proteins (generated during electroformation of GUVs). These ‘naturally occurring’ ILVs were used as a benchmark for the typical average membrane composition of the parent GUV. Therefore a higher Rh-PE fluorescence than this benchmark would indicate ESCRT-induced ILVs are enriched in L_d_ phase and a lower average Rh-PE fluorescence would indicate that ESCRT-induced ILVs are enriched in L_o_ phase.

## Acknowledgements

This work was funded by the UK Engineering and Physical Sciences Research Council (EPSRC): EP/M027929/1 (PAB) and EP/M027821/1 (BC). We also thank Dr. Daniel Mitchell for expression and purification of the Vps4 subunit.

